# Delayed SARS-CoV-2 Spread and Olfactory Cell Lineage Impairment in Close-Contact Infection Syrian Hamster Models

**DOI:** 10.1101/2022.09.03.506499

**Authors:** Rumi Ueha, Toshihiro Ito, Satoshi Ueha, Ryutaro Furukawa, Masahiro Kitabatake, Noriko Ouji-Sageshima, Tsukasa Uranaka, Hirotaka Tanaka, Hironobu Nishijima, Kenji Kondo, Tatsuya Yamasoba

## Abstract

**Objectives:** Close contact with patients with COVID-19 is speculated to be the most common cause of viral transmission, but the pathogenesis of COVID-19 by close contact remains to be elucidated. In addition, despite olfactory impairment being a unique complication of COVID-19, the impact of SARS-CoV-2 on the olfactory cell lineage has not been fully validated. This study aimed to elucidate close-contact viral transmission to the nose and lungs and to investigate the temporal damage in the olfactory receptor neuron (ORN) lineage caused by SARS-CoV-2.

**Methods:** Syrian hamsters were orally administered SARS-CoV-2 as direct-infection models. On day 7 after inoculation, infected and uninfected hamsters were housed in the same cage for 30 minutes. These uninfected hamsters were subsequently assigned to a close-contact group. First, viral presence in the nose and lungs was verified in the infection and close-contact groups at several time points. Next, the impacts on the olfactory epithelium, including olfactory progenitors, immature ORNs, and mature ORNs, were examined histologically. Then, the viral transmission status and chronological changes in tissue damage were compared between the direct-infection and close-contact groups.

**Results:** In the close-contact group, viral presence could not be detected in both the nose and lungs on day 3, and the virus was identified in both tissues on day 7. In the direct-infection group, the viral load was highest in the nose and lungs on day 3, decreased on day 7, and was no longer detectable on day 14. Histologically, in the direct-infection group, mature ORNs were most depleted on day 3 (p < 0.001) and showed a recovery trend on day 14, with similar trends for olfactory progenitors and immature ORNs. In the close-contact group, there was no obvious tissue damage on day 3, but on day 7, the number of all ORN lineage cells significantly decreased (p < 0.001).

**Conclusion:** SARS-CoV-2 was transmitted even after brief contact and subsequent olfactory epithelium and lung damage occurred more than 3 days after the trigger of infection. The present study also indicated that SARS-CoV-2 damages all ORN lineage cells, but this damage can begin to recover approximately 14 days post infection.

## 1 Introduction

Coronavirus disease (COVID-19), caused by severe acute respiratory syndrome coronavirus 2 (SARS-CoV-2), is a pandemic since the end of December 2019 (1, 2) that remains to be contained; rather, the infected population is increasing worldwide. The incubation period from SARS-CoV-2 exposure to the onset of COVID-19 symptoms is reported to be approximately 6 days, and the viral load in droplets increases several days prior to the onset of symptoms in patients with COVID-19 (3-5). Accordingly, if virus carriers do not take appropriate infection control measures during the asymptomatic period, they may play an important role in unintentional COVID-19 spread (6, 7). The most likely asymptomatic carriers are considered close contacts, defined as individuals who had contact with infected persons for more than 15 minutes within a 1-m distance without properly wearing masks, and these individuals are considered at high risk of exposure to SARS-CoV-2 and developing infection (8, 9). Nevertheless, how the virus spreads and multiplies in the body after brief close contact with an infected person has not been sufficiently studied.

The nasal cavity comprises important tissue for the replication of SARS-CoV-2, and SARS-CoV-2 can cause chemosensory dysfunction and affect olfaction (10, 11). In the early days of the COVID-19 pandemic, olfactory dysfunction was often the first manifestation of COVID-19 (12). The prevalence and severity of COVID-19-related olfactory dysfunction has decreased since the omicron variant became prevalent, but it remains an important issue as a sequela of COVID-19 (13). Despite the nose being an essential sensory organ, it has not been fully elucidated whether short-term close contact with an infected person can cause nasal damage. In previous studies using animal models, animals were administered high doses of SARS-CoV-2 and severe nasal epithelial damage was identified within a few days after viral exposure (14-17). However, as high doses of viruses are unlikely to be taken in at once in a real-life environment, the damage to the olfactory epithelium (OE) of the nose in these SARS-CoV-2 infection models may be discrepant from the actual situation. Therefore, it is imperative to validate close-contact infection models that readily resemble the infection situation in daily life. In addition, SARS-CoV-2 can be transmitted both nasally and orally in daily life, and the longitudinal tissue damage after nasal and oral virus administration needs to be evaluated.

The OE is composed of supporting cells (sustentacular cells), basal progenitor cells, immature olfactory receptor cells (ORNs), and mature ORNs (18). Basal cells and supporting cells are particularly susceptible to damage by SARS-CoV-2 infection, and the entire OE may be denuded depending on the site (10, 14, 15, 19). This OE impairment improves over time, and the OE is nearly normalized within approximately 1 month after SARS-CoV-2 infection, although the regenerative kinetics may differ according to the nose area, such as the nasal septum and lateral nasal turbinate (14). Although temporal histological changes in the epithelial thickness of the OE over time have been reported, the impact of SARS-CoV-2 on the various cell groups comprising the OE over time has not been verified. Furthermore, as the susceptibility to damage and regeneration progression varies depending on the location in the nose (20, 21), it would be meaningful to examine the effects of SARS-CoV-2 on the OE, especially on the ORN lineage, in a site-specific manner.

In this study, to elucidate the biological effects of short-term close contact with SARS-CoV-2 infected individuals, we created short-term contact models and examined histological and molecular biological changes over time in the nose and lungs. Next, we examined the temporal changes in the OE in a nasal-cavity site-specific manner in hamsters infected with SARS-CoV-2. Last, we compared the time course of virus spread in tissues between direct-infection and close-contact models.

## 2 Materials and Methods

### 2.1 Animals

We selected Syrian hamsters (*Mesocricetus auratus*) as the animal model for this study because Syrian golden hamsters have been used in various studies related to SARS-CoV-2 and are recognized as an excellent small animal model for SARS-CoV-2 infection (22, 23). Six-week-old, male Syrian hamsters were purchased from Japan SLC (Hamamatsu, Shizuoka, Japan) and maintained in a specific pathogen-free environment at the Animal Research Center of Nara Medical University. The animal experimental protocols followed the ARRIVE guidelines and were approved by The Animal Care and Use Committee at Nara Medical University (approval number, 12922). All procedures were performed in compliance with relevant guidelines on the Care and Use of Laboratory Animals, Nara Medical University and Animal Research, and Animal Care and Use Committee of the University of Tokyo.

### 2.2 Animal model preparation

SARS-CoV-2 infected animal models were prepared following previously reported methods (19, 24). In short, the SARS-CoV-2 strain (JPN/TY/WK-521, provided by the National Institute of Infectious Diseases, Japan) was used. Twenty-four hamsters were distributed into three groups: four hamsters in the control group, 12 hamsters in the SARS-CoV-2 transoral infection group (infected group), and eight hamsters in the short-term close-contact group (contact group). All experiments using SARS-CoV-2 were performed at the biosafety level 3 experimental facility of Nara Medical University, Japan. The time course is shown in Figure 1A.

**Figure 1.**
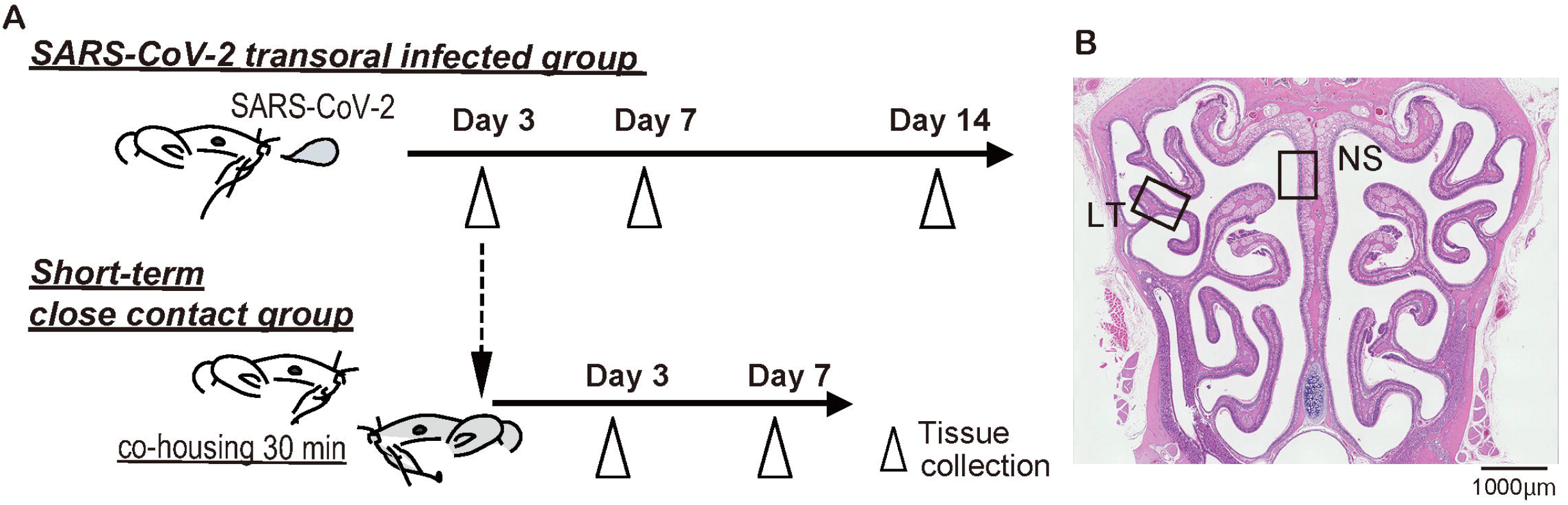
Experimental timeline and nasal structure. **A**: Hamsters were administered orally severe acute respiratory syndrome coronavirus 2 (SARS-CoV-2) (1.0×10^3^ pfu). To prepare the short-term close-contact models, the uninfected hamsters were cohoused for 30 minutes with SARS-CoV-2 infected hamsters (3 days after SARS-CoV-2 inoculation), and then separated. **B**: Representative images of the olfactory epithelium from control hamsters. The boxes indicate the regions of the dorsal nasal septum (NS) and lateral turbinate (LT) areas.

The hamsters were anesthetized with an intraperitoneal injection of pentobarbital (10 mg/mL, 0.8 mL/hamster) before virus inoculation. To the SARS-CoV-2 transoral infection group, 50 μL of virus solution diluted with saline (including 1.0×10^3^ pfu of SARS-CoV-2) was administered orally as previously reported (19, 24). To the control group, 50 μL of saline was administered. At 3, 7, and 14 days after inoculation, the SARS-CoV-2-inoculated hamsters were euthanized by intraperitoneal injection of 1.0 ml sodium pentobarbital (10 mg/mL) followed by cardiac exsanguination.

To prepare the short-term close-contact models, uninfected hamsters were transferred to the cages of infected hamster on day 3 of SARS-CoV-2 administration and cohoused. After 30 minutes of cohabitation, the short-term close-contact hamsters and directly infected hamsters were separated. Three and 7 days after contact, the close-contact-group hamsters were euthanized. The noses were sampled for histopathological examination, and the lungs were sampled for histopathological examination and quantitative polymerase chain reaction (qPCR).

### 2.3 RNA extraction and RT-qPCR

We validated SARS-CoV-2 viral RNA in the lungs to confirm the SARS-CoV-2 infection status of each hamster. Total RNA was isolated from the lung using NucleoSpin^®^ RNA (Macherey-Nagel, Düren, Germany), and then converted to cDNA using a High-Capacity cDNA Reverse Transcription Kit (Thermo Fisher Scientific, Waltham, MA, USA), according to the previous protocol (19, 24) and the manufacturer’s instructions. RT-qPCR analysis was performed using a StepOnePlus Real-Time PCR System (Thermo Fisher Scientific). The gene-specific primers and probes used were: *Gapdh* as endogenous control (TaqMan assay Cg04424038) and SARS-CoV-2 nucleocapsid gene (forward: 5’-AAATTTTGGGGACCAGGAAC −3’, reverse: 5’-TGGCAGCTGTGTAGGTCAAC −3’, the TaqMan probe: FAM-ATGTCGCGCATTGGCATGGA-BHQ). The expression levels of each gene were normalized to the level of *Gapdh* expression for each sample.

### 2.4 Tissue preparation

Immediately after tissue harvesting, the lungs and nasal tissues assigned for histological analyses were gently irrigated and fixed in 4% paraformaldehyde for 14 days. Then, the tissue samples were decalcified, dehydrated in graded ethanol solutions, and embedded in paraffin. Coronal sections were obtained from all samples at the level of the anterior end of the olfactory bulb for histological analysis of the olfactory mucosa (18, 25-27). Four-micrometer-thick paraffin sections were deparaffinized in xylene and rehydrated in ethanol before staining.

### 2.5 Histological analyses

Hematoxylin and eosin staining was performed to evaluate the overall tissue structure. For immunostaining, deparaffinized sections were treated with 3% hydrogen peroxide to block endogenous peroxidase activity and then incubated in Blocking One solution (Nacalai Tesque, Kyoto, Japan) to block non-specific immunoglobulin binding. After antigen retrieval, the samples were incubated with primary antibodies, followed by secondary antibodies.

The primary antibodies used in this study are listed in Table 1. Anti-SARS-CoV-2 nucleocapsid antibody was used to identify SARS-CoV-2. The following antibodies were used to evaluate ORN neurogenesis: sex-determining region Y-box 2 (SOX2), expressed by proliferating stem cells or progenitor cells in the basal layer of the OE; growth-associated protein 43 (GAP43), expressed by immature ORNs in the OE; and olfactory marker protein (OMP), expressed by mature ORNs in the OE.

**Table 1.** The primary antibodies used in this study

| Primary antibody | Source | Catalog No. | Host | Type | Dilution |
| --- | --- | --- | --- | --- | --- |
| SARS-CoV-2<br>nucleocapsid | GeneTex (Irvine, CA, USA) | GTX135357 | Rabbit | polyclonal | 1:1000 |
| SOX2 | Abcam (Cambridge, UK), | ab92494 | Rabbit | monoclonal | 1:300 |
| GAP43 | Novus Biologicals (Centennial, CO, USA) | NB300-143B | Rabbit | polyclonal | 1:1000 |
| OMP | Wako (Tokyo, Japan) | 019-22291 | Goat | polyclonal | 1:8000 |

The ORNs are classified into four groups according to their zonal-expression patterns, and odorant receptors are expressed by sensory neurons distributed within one of four circumscribed zones (28-30). Of these, zone 1 is determined by co-localization with NQO1 expression and zones 2–4 are determined by OCAM expression (28-31). Our previous studies have shown that the OE of the dorsal nasal septum area represents OCAM^-^/NQO1^+^ expression (Zone 1) and that of the lateral area represents OCAM^+^/NQO1^-^ expression (zones 2–4). Therefore, to analyze the OE, we divided coronal sections of the OE into two areas along the zonal organization: the dorsal nasal septum (NS) area and lateral turbinate (LT) area (Figure 1B).

Images were captured using a digital microscope camera (Keyence BZ-X700, Osaka, Japan) with 10× and 40× objective lenses. OMP^+^ ORNs, SOX2^+^ ORN progenitors, and GAP43^+^ immature ORNs in a 300-μm region of each area were counted in the right and left sides of each sample. The number of each cell type was quantitatively analyzed using sections with immunostaining for each antigen and counterstaining with hematoxylin.

### 2.6 Statistical analysis

Statistical comparisons between groups were performed with one-way analysis of variance using GraphPad Prism software (version 6.7; GraphPad Software, Inc., San Diego, CA, USA, www.graphpad.com). qPCR data were subjected to logarithmic transformation prior to analysis. Results with *p* < 0.05 were considered statistically significant.

## 3 Results

### 3.1 Time course of SARS-CoV-2 infection after oral virus inoculation is analogous in the nose and lungs with a tendency to subside 7 days post infection

To confirm that SARS-CoV-2 infection was established, we preliminary evaluated the presence of the virus in the noses and lungs of the SARS-CoV-2-infected hamsters on day 3 after oral inoculation with immunohistochemistry and RT-qPCR. The virus was identified in the noses with immunohistochemistry and in the lungs with immunohistochemistry and RT-qPCR. Viruses were found in various areas of the nasal cavity but were not evenly spread throughout the nasal mucosa. On day 7 post infection, the presence of virus diminished in both the noses and lungs, and the viral load in the lungs was reduced to less than 1 in 1000 according to the qPCR analyses. In the nasal cavity, the virus was rarely observed in the area near the nasal septum, but the virus was still present in the outer region. On the 14th day of infection, no virus was identified histologically in either the noses or lungs (Figure 2A, 2B) and no viral genes could be detected with qPCR (Figure 2C).

**Figure 2.**
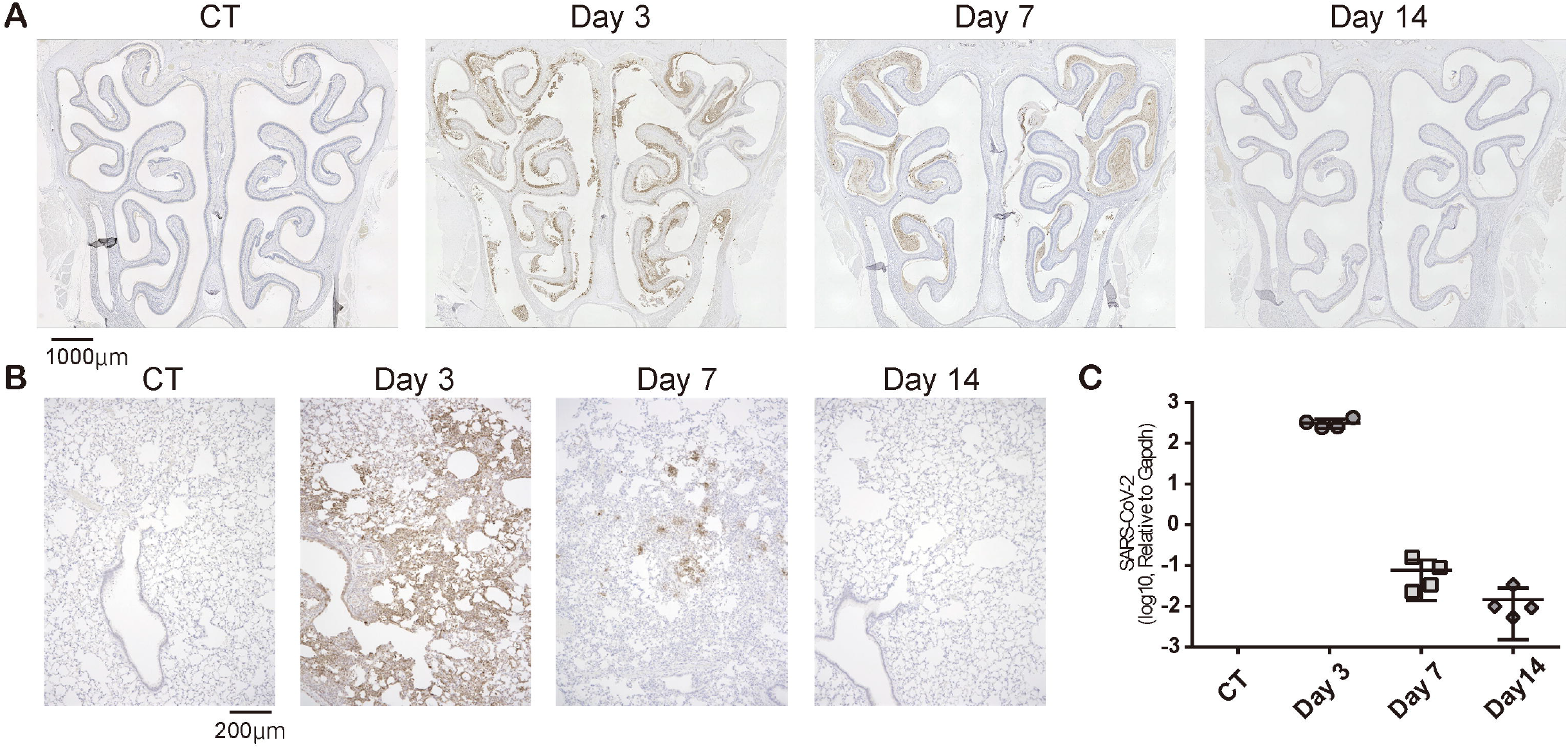
Temporal SARS-CoV-2 infection findings in the nose and lungs. **A**: Representative images of immunohistological staining of severe acute respiratory syndrome coronavirus 2 (SARS-CoV-2) in control (CT) hamsters and SARS-CoV-2 hamsters on days 3, 7, and 14. SARS-CoV-2 is shown in brown. **B**: Immunohistochemistry staining of SARS-CoV-2 in the lung. **C**: SARS-CoV-2 gene detection with RT-quantitative polymerase chain reaction in the CT hamster and SARS-CoV-2 hamsters on days 3, 7, and 14. SARS-CoV-2: severe acute respiratory syndrome coronavirus 2

Taken together, in the virus-direct inoculation model, the infection findings were severe on approximately day 3, and the viral load decreased on day 7 after virus administration; by day 14, the virus had almost completely cleared from the nose and lungs (Figure 2).

### 3.2 No SARS-CoV-2 infection signs appeared in the first few days after short-term close contact but became apparent after a certain period following contact

Next, we examined whether SARS-CoV-2 could infect the nose and lungs over time by brief contact with infected animals. SARS-CoV-2 was not identified in the noses or lungs on day 3 after short-term contact, but the virus was extensively identified in both the nose and lungs of all hamsters on day 7 after close contact. In the nasal cavity, the virus was more apparent in the LT area than in the NS area. As for the lungs, on day 7, more virus was present in the close-contact group than in the direct-infection group, at an almost similar level to that of day 3 in the direct-infection group (Figure 3). The period for the virus to be identified in the tissues was markedly delayed compared to the direct-infection group.

**Figure 3.**
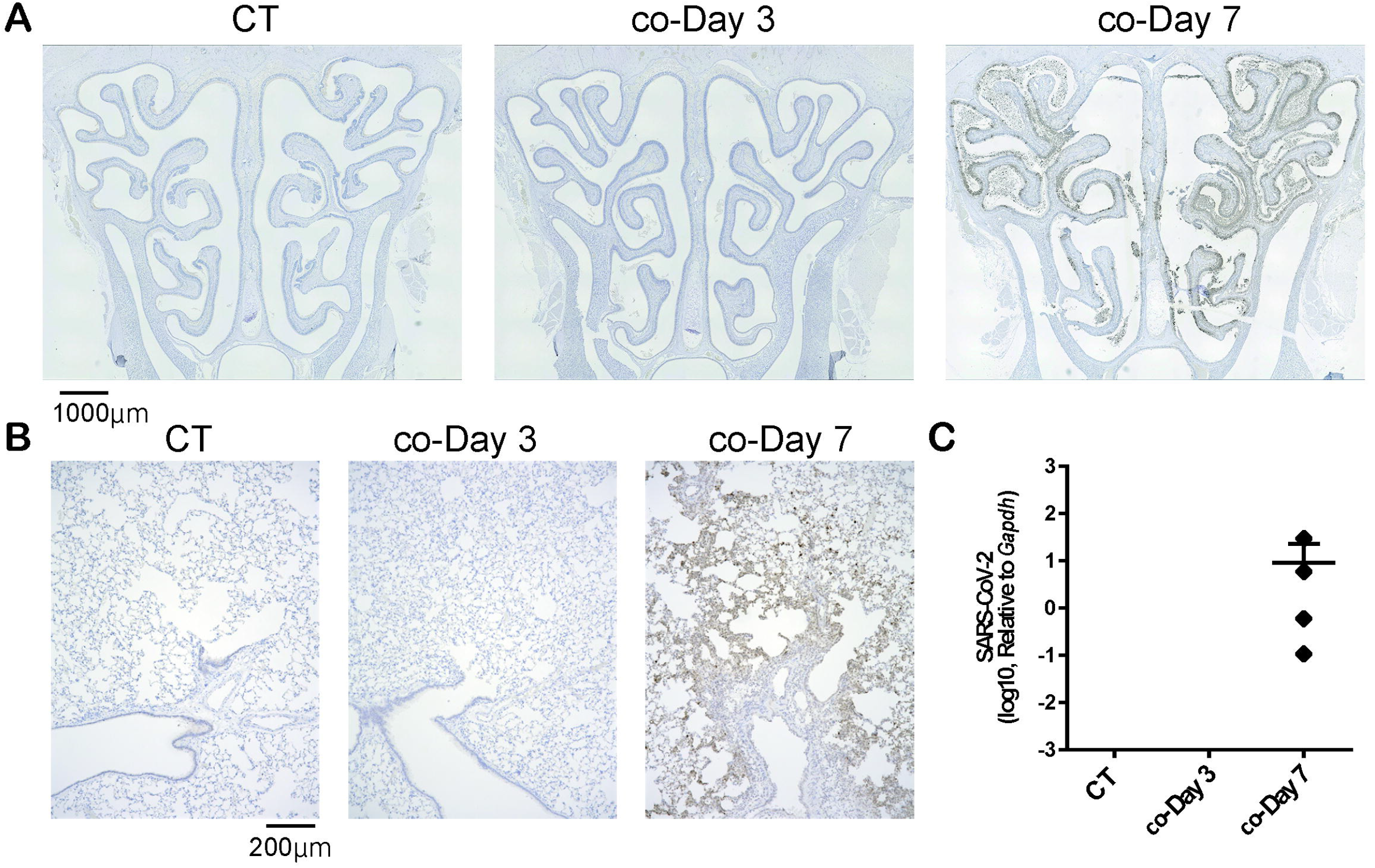
Representative SARS-CoV-2 infection findings in the nose and lungs after short-term close contact with SARS-CoV-2 infected animals. **A, B**: Representative images of SARS-CoV-2 staining in the nose (**A**) and lungs (**B**) 3 and 7 days after short-term close contact with SARS-CoV-2 infected animals (co-Day 3, co-Day 7). The number of each type of cell in a 300-μm region of each area was counted in the right and left sides of each sample (n = 4, each group). **C**: SARS-CoV-2 gene detection with RT-quantitative polymerase chain reaction in the CT hamster and contact hamsters on days 3 and 7. SARS-CoV-2: severe acute respiratory syndrome coronavirus 2

### 3.3 Transoral SARS-CoV-2 infection influences the ORNs at various differentiation stages over time

Subsequently, we examined the temporal effects of SARS-CoV-2 infection on the ORN lineage in the direct-infection model. We found differences in tissue damage between the NS and LT areas; in the NS area, on day 3, almost all superficial cells above the basal layer were missing, especially in the severely damaged areas, but the thickness of the OE tended to recover on day 14 of infection, suggesting OE regeneration. Conversely, in the LT area, the OE did not detach from the basal layer during days 3 to 14 after virus administration (Figure 4A). Figure 4 illustrates representative findings in the most severely affected areas of the OE.

**Figure 4.**
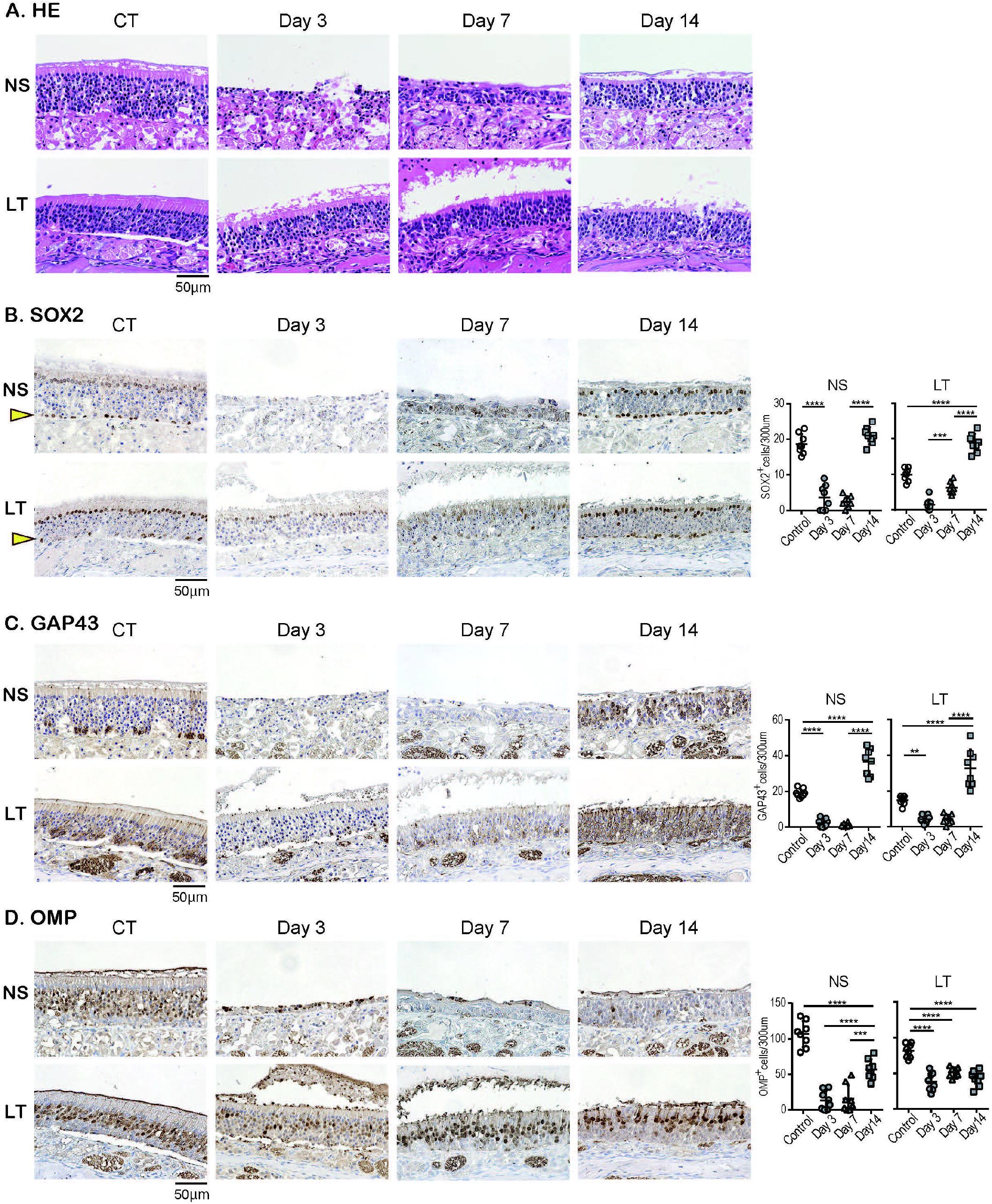
Effects of SARS-CoV-2 infection on the olfactory receptor neuron lineage. **A**: Representative hematoxylin and eosin staining (HE) images of the olfactory epithelium in control (CT) hamsters and SARS-CoV-2 hamsters on days 3, 7, and 14. **B-D**: Representative images of immunohistological staining in CT and SARS-CoV-2 hamsters. Nasal septum (NS) area and lateral turbinate (LT) area are shown in magnified view. Sex-determining region Y-box 2 (SOX2)^+^ progenitor cells (**B**), growth-associated protein 4 (GAP43)^+^ immature olfactory receptor neurons (ORNs) (**C**), and olfactory marker protein (OMP)^+^ ORNs (**D**) are shown in brown. The basal layer is indicated by arrows. Tissue sections were counterstained with the nuclear dye hematoxylin (blue). Numbers of SOX2^+^ ORN progenitors per 300 μm of the basal layer, and GAP43^+^ immature ORNs and OMP^+^ mature ORNs per 300 μm of olfactory epithelium in each area are counted in CT or SARS-CoV-2 hamsters. **, *P* < 0.01; ***, *P* < 0.001; ****, *P* < 0.0001. SARS-CoV-2: severe acute respiratory syndrome coronavirus 2

In addition, we examined the impact of SARS-CoV-2 on all ORN lineage cells depending on the OE area. SOX2^+^ olfactory progenitors were scarcely identified in the NS area on days 3 and 7 but recovered to the same level as that in the control group on day 14. In contrast, in the LT area, the number of SOX2^+^ cells decreased on days 3 and 7, but it became higher on day 14 compared to that of the control group (Figure 4B). GAP43^+^ immature ORNs could scarcely be seen in both the NS and LT areas on days 3 and 7, whereas the number of these cells increased on day 14 compared to the control group (Figure 4C). OMP^+^ ORNs could rarely be observed in the NS area on days 3 and 7, but a recovery tendency in cell counts was evident on day 14, but did not reach the level in the control group. In the LT area, OMR^+^ ORNs existed to some extent in the OE on days 3, 7, and 14, although they were fewer than those in the control group (Figure 4D). Thus, the ORN lineage was differently affected by SARS-CoV-2 infection in the NS and LT areas.

### 3.4 Short-term close contact with SARS-CoV-2 infected models causes late damage in all ORN lineage cells

Last, we investigated how late-onset SARS-CoV-2 infection by short-term close contact with infected animals influences the OE over time. Although no structural changes in the OE were observed on day 3 after short-term close contact, marked disruption of the OE in the NS area was observed on day 7 (Figure 5A). Immunohistological examination showed no significant differences in the number of all ORN lineage cells of the OE on the 3rd day after contact compared with the control group, which supported the observation that no virus was identified in the tissue. Conversely, in the OE on day 7 after contact, neither SOX2^+^ olfactory progenitors nor GAP43^+^ immature ORNs could be identified in the NS or LT area, with only a few OMP^+^ mature ORNs in the LT area (Figure 5B-D). Thus, it was evident that short-term close contact could impair the ORN lineage, and the timing of this impairment was later than that by direct virus administration.

**Figure 5.**
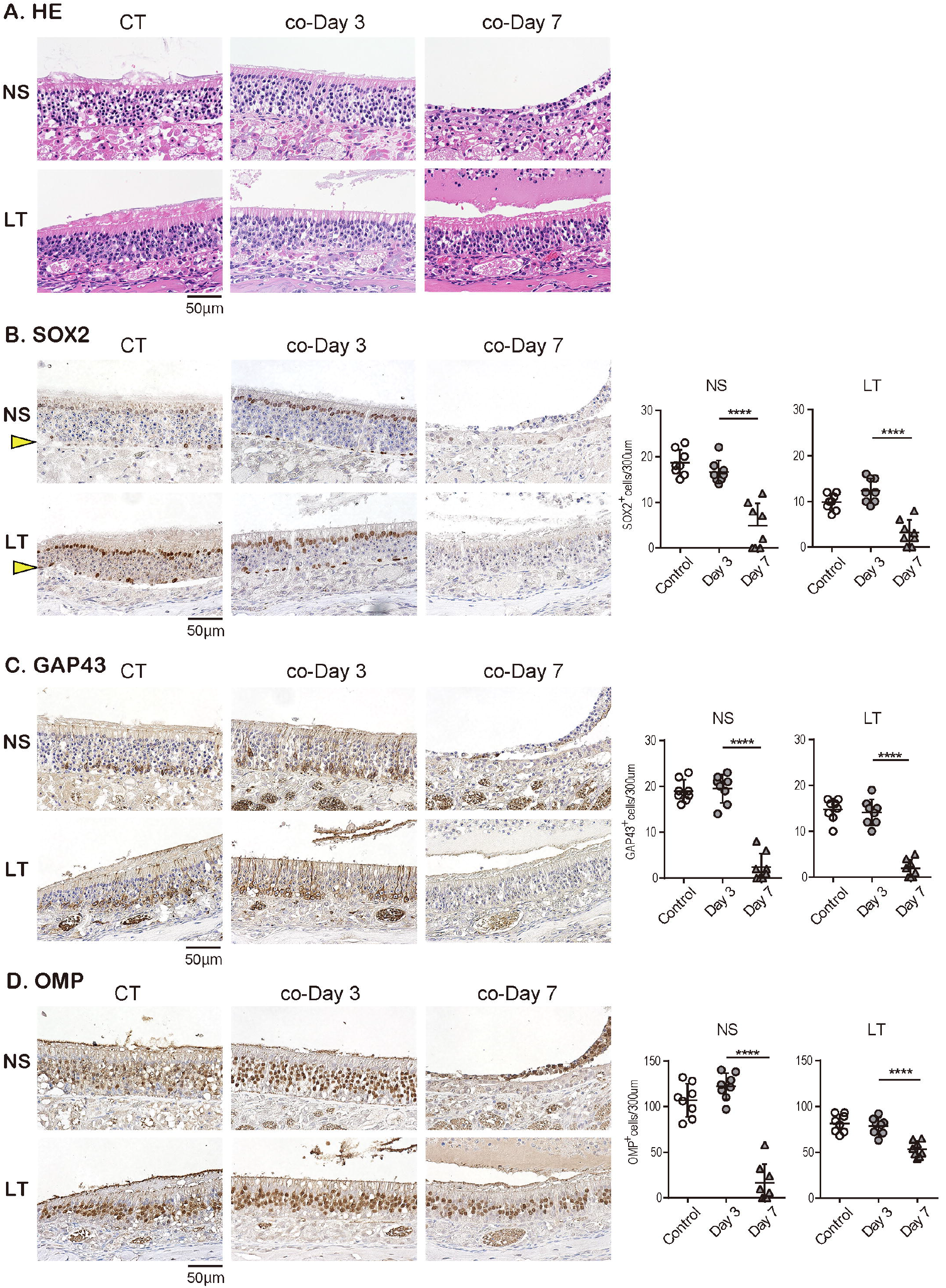
Effects of SARS-CoV-2 on the olfactory receptor neuron lineage after short-term close contact. **A**: Representative hematoxylin and eosin staining (HE) images of the olfactory epithelium in control (CT) hamsters and contact hamsters on days 3 and 7 (co-Day 3, co-Day7). **B–D**: Representative images of immunohistological staining. The nasal septum (NS) area and lateral turbinate (LT) area are shown in magnified view. Sex-determining region Y-box 2 (SOX2)^+^ progenitor cells (**B**), growth-associated protein 4 (GAP43)^+^ immature olfactory receptor neurons (ORNs) (**C**), and olfactory marker protein (OMP)^+^ ORNs (**D**) are shown in brown. The basal layer is indicated by arrows. Tissue sections were counterstained with the nuclear dye hematoxylin (blue). Numbers of SOX2^+^ ORN progenitors per 300 μm of the basal layer and GAP43^+^ immature ORNs and OMP^+^ mature ORNs per 300 μm of olfactory epithelium in each area are counted in CT or contact hamsters. ****, *P* < 0.0001. SARS-CoV-2: severe acute respiratory syndrome coronavirus 2

## 4 Discussion

The present study showed that short-term close contact with infected hamsters did not cause nasal or pulmonary damage by day 3 but resulted in widespread infection of the nose and lungs within 7 days. In contrast, direct SARS-CoV-2 administration caused tissue damage in the nose and lungs with the virus being detected within a few days. Thereafter, the viral load in the tissues decreased over time, and no virus was identified in the nose or lungs 14 days post infection. We also demonstrated that SARS-CoV-2 extensively damaged the OE, and the degree of OE damage over time varied depending on the OE site. The numbers of ORN-related cells were reduced in all lineage cells with time and then tended to recover (Figure 6).

**Figure 6.**
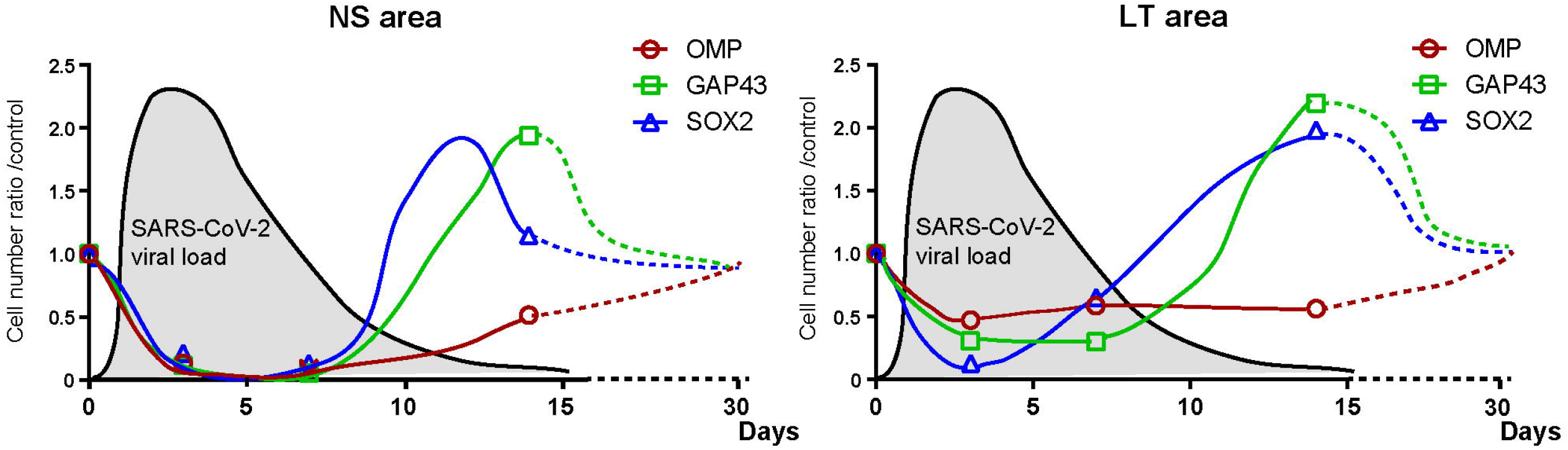
Hypothesis of temporal changes in the SARS-CoV-2 viral load in the lungs and the number of olfactory receptor neuron-related cells in different nasal areas Changes over time in the ratio of the number of olfactory receptor neuron lineage cells divided by that of the control group at each time point are displayed in the graph. The dashed line shows the speculated future trend. In addition, the estimated viral load of SARS-CoV-2 in the lungs is overlaid on each graph. NS: nasal septum, LT: lateral turbinate, OMP: OMP^+^ mature olfactory receptor neurons, GAP43: GAP43^+^ immature olfactory receptor neurons, SARS-CoV-2: severe acute respiratory syndrome coronavirus 2, SOX2: SOX2^+^ olfactory progenitors.

Clinically, the incubation period, which required for COVID-19 symptoms to appear, is approximately 6 days (3, 32), and SARS-CoV-2 RNA shedding lasts approximately 17 days in the upper respiratory tract (33, 34). Animal studies using SARS-CoV-2 infection models have reported that the viral load is highest on days 2–5 after virus administration (14, 35, 36) and the virus is eliminated from the tissues on days 8–10 after administration (14, 35); the direct-infection model in the present study showed a similar trend in viral load. In this study, the SARS-CoV-2 viral load in hamsters increased after 1 week, despite a short contact time of 30 minutes. Therefore, there is a risk of viral transmission when in contact with SARS-CoV-2 infected individuals, even if it is only for a short period of time (approximately 30 minutes). Even though the virus could not be identified in the nose or lungs on the 3rd day after close contact, severe damage in both tissues was observed on the 7th day after contact, suggesting that the 4th to 6th day after contact with the infected animal was the time when the virus, which had remained latent in the meantime, multiplied. Although the present validation could only examine a period of 7 days after close contact, on day 7, the viral load in the lungs of the animals in the close-contact group was higher than that of the animals in the direct-infection group, implying that another week or longer may be required for the viral load level to become sufficiently low. Temporal changes in viral load and tissue damage after close contact remain a future topic of investigation.

Regarding the transmission form in this short-term close-contact model, as the mouths and noses of the hamsters were not covered, various routes of infection may be considered, including contact, droplet, and airborne infection. Accordingly, if someone infected with SARS-CoV-2 talks or eats with others without a mask for 30 minutes, he or she can spread the infection to others. The percentage of people contracting SARS-CoV-2 and being asymptomatic is estimated to range from 1.2 to 12.9% (37, 38). It has now been more than 2 years since COVID-19 became endemic, and the spread of infection has not been well-controlled. If asymptomatic carriers do not take appropriate infection control measures, they may spread the SARS-CoV-2 infection without realizing it.

Olfactory dysfunction caused by SARS-CoV-2 is quite common, and it is now well established how the virus spreads to various olfactory-related tissues, such as the olfactory mucosa, olfactory bulb, and olfactory cortex (39-41). Regarding the effects of SARS-CoV-2 on the OE, it has been reported that the degree of epithelial damage and recovery rate of the OE differ depending on the location in the nasal cavity (14). However, the impact of SARS-CoV-2 on each ORN lineage cell has not been sufficiently verified. We previously reported that all ORN lineage cells are impaired by SARS-CoV-2 (19), and in this study, we verified related longitudinal effects for the first time to our knowledge.

In the NS area, SARS-CoV-2 infection caused shedding of almost all cells of the OE and greatly thinned the OE. The supporting cells of the OE express high levels of ACE2 (15, 42, 43), which may be susceptible to infection by SARS-CoV-2. It is possible that SARS-CoV-2 infection of the supporting cells interferes with their structural support, resulting in widespread epithelial shedding. ACE2 is also expressed in the basal olfactory progenitor cells (10, 42), and the low number of SOX2-positive cells on day 3 post infection suggests that SARS-CoV-2 infection may have directly impaired the olfactory progenitor cells. In addition, as the other receptor of SARS-CoV-2, neuropilin-1 is expressed in almost all cells of the OE (44, 45), it is not surprising that all ORN lineage cells could be affected by SARS-CoV-2. Thereafter, by day 14, the olfactory progenitors and immature ORNs had developed, and epithelial regeneration had become active, but mature ORNs had not sufficiently regenerated, suggesting that overall recovery from SARS-CoV-2 infection-induced OE damage may require longer than 14 days (Figure 6). In fact, Urata et al. (14) and Reyna et al. (35) reported that in the NS, more than 21 days are needed for the OE thickness to recover to normal. Based on the results of this validation and previous reports, our hypothesis regarding the temporal impact of SARS-CoV-2 on the ORN lineage is presented in Figure 6.

In the LT area, the number of OMP-positive cells did not considerably change over time, possibly because some ORNs are impaired by the virus, whereas the unimpaired ORNs remain present in the OE. Thus, it is conceivable that the SARS-CoV-2 receptor expression of the cells in that area may differ depending on the olfactory epithelium site. Moreover, given the increase in GAP43^+^ immature ORNs in the LT area on day 14, the number of impaired OMP^+^ mature ORNs is expected to increase on day 14 or later. As shown in Figure 6, the recovery of mature OMPs may need longer in the LT area than in the NS area because of the later timing of the numerical peak of ORN progenitors and immature ORNs in the LT area than in the NS area. Future studies are needed for further observations of the long-term infectious course after short-term close contact.

In conclusion, the present study demonstrated that SARS-CoV-2 can be transmitted even after brief contact and that subsequent OE damage occurs more than 3 days after the trigger of infection. Moreover, SARS-CoV-2 could damage the olfactory receptor system, but the damage could begin to recover approximately 14 days post infection. For SARS-CoV-2 infection control, it is desirable to have a global discussion on the infection control measures that should be implemented and to create common worldwide rules.

## 6 Authors’ contributions

RU developed the concept, designed and performed the experiments, analyzed the data, produced the figures, and wrote the initial draft of the manuscript. TI developed the concept, prepared the animal models, performed some of the experiments, and analyzed the data. SU developed the concept, designed the experiments, and revised the manuscript. RF, MK, and NOS prepared the animal models, performed some of the experiments, and analyzed the data. TU, HT, and HN performed some of the experiments and analyzed the data. KK and TY developed the concept and critically revised the manuscript. All authors contributed to interpretation of the data and writing of the manuscript.

## 8 Acknowledgements

Not applicable.

## Availability of data and materials

The datasets used and/or analyzed during the current study are available from the corresponding author on reasonable request.

